# The Benefits of Mycorrhizae are Frequency-Dependent: A Case Study With a Non-Mycorrhizal Mutant of *Pisum Sativum*

**DOI:** 10.1101/2020.09.08.288084

**Authors:** Gordon G. McNickle, Frédérique C. Guinel, Anastasia E. Sniderhan, Cory A. Wallace, Allison S. McManus, Melissa M. Fafard, Jennifer L Baltzer

**Affiliations:** Department of Botany and Plant Pathology, Purdue University, 915 W State St., West Lafayette, IN, 47907; Purdue Center for Plant Biology, Purdue University, West Lafayette, IN, 47907; Wilfrid Laurier University, Department of Biology, 75 University Avenue West, Waterloo, ON N2L 3C5

## Abstract

1. Mutualisms are remarkably common in the plant kingdom. The mycorrhizal association which involves plant roots and soil fungi is particularly common, and found among members of the majority of plant families. This association is a resource-resource mutualism, where plants trade carbon-based compounds for nutrients, such as phosphorus and nitrogen, mined by the fungi.
2. Evolutionary models usually assume that a mutation grants a small number of individual plants the ability to associate with mycorrhizal fungi, and that this subsequently spreads through the population resulting in the evolution of mutualism. This frequency-dependent hypothesis has been difficult to test, because it is rare to have members of the same species that are capable and incapable of forming the mutualism.
3. Here we describe the results of an experiment that took advantage of a mutant pea (*Pisum sativum* L. R25) that is incapable of forming mycorrhizal (or rhizobial) associations, and differs from the wildtype (*P. sativum* cv. Sparkle) by a single recessive Mendelian allele (*Pssym8*). We grew each genotype either alone or in every combination of pairwise mixed- or same-genotype. We also present an evolutionary matrix game, which we parameterize from the experimental ^15^N results, that allows us to estimate the costs and benefits of the mutualism.
4. We find that there was no difference between R25 and WT when grown with a competitor of the same genotype, but when R25 and WT compete, WT has a significant fitness advantage. From the model, we estimate that the benefit in units of fitness (g pod mass) obtained from direct plant nitrogen uptake is 22.2 g, and mycorrhizae increase this by only 0.6 g. The costs of plant nitrogen uptake are 9.4 g, while the cost of trade with mycorrhizae is 0.1g.
5. From the model and experiment, we conclude that this relatively small cost-benefit ratio of the mycorrhizal association is enough to drive the evolution of mutualism in frequency-dependent selection. However, without the mutant R25 genotype we would not have been able to draw this conclusion. This validation of frequency-dependent evolutionary models is important for continued theoretical development.

## INTRODUCTION

Mutualisms are remarkably common within the plant kingdom. For example, estimates suggest that 80% of terrestrial plant species (92% of families) associate with soil fungi to form a mutualistic mycorrhizal association (Wang & Qiu 2006). The mycorrhizal association is a resource-trading mutualism where plants exchange carbon-based compounds for mineral nutrients with fungi. Historically, it was thought that mycorrhizae primarily provided plants with phosphorus (Sanders & Tinker 1973), but recent work has shown that arbuscular mycorrhizal fungi also transfer substantial amounts of nitrogen (N) to plants from decomposed organic material (Leigh, Hodge & Fitter 2009; Hodge & Fitter 2010). Nitrogen is generally the main limiting resource for terrestrial plants, and thus obtaining N via mutualism has the potential to dramatically affect plant fitness (Chapin, Vitousek & Vancleve 1986; Vitousek & Howarth 1991).

Indeed, analysis of evolutionary models suggests that mutualisms such as mycorrhizae evolve when there is a fitness advantage to individuals that engage in cooperative mutualisms relative to those individuals that do not engage in mutualism (Nash 1950; Axelrod & Hamilton 1981; Denison *et al*. 2003). Though the details differ, these eco-evolutionary models begin with an assumption that the ancestral population had no mutualism, then an individual undergoes some mutation(s) that give it access to a mutualistic partner. While mutation is necessary for evolution by natural selection, on its own, mutation is not sufficient to cause a stable evolutionary change. Once mutation confers the mutualism ability, those mutant individuals must subsequently out-compete non-mutualistic residents in the population, resulting in the mutualism spreading through the population. While this is logically intuitive, and supported by a wide array of models, testing this preposition is difficult because it requires a mixture of individuals from the same species that can and cannot form a mutualistic association.

It is straightforward to assess competition among plants which are either in a mutualistic relationship or are not through the use of some combination of soil sterilization or inoculation experiments (Fitter 1977; Allen & Allen 1984; Moora & Zobel 1996; Hodge *et al*. 2000; Hodge 2001; Hodge 2003). In general, these studies find that plant-plant competition is stronger when plants associate with mycorrhizae than when they do not (Fitter 1977; Hodge 2003), or that there is no difference among competing plants with or without mycorrhizae (Allen & Allen 1984; Moora & Zobel 1996). At first glance, this seems to call into question the evolutionary models of mutualism. However, these studies assume static relationships between the strategy (e.g. mutualism vs no mutualism) and fitness of an individual regardless of the strategies of its neighbours, thus neglecting tests of frequency-dependent selection. The key experimental treatment required to examine the frequency-dependent evolution of mutualism, which would contain a mixture of mycorrhizal and non-mycorrhizal competitors from the same population, was absent from all of these studies because this situation is typically not possible to experimentally create.

Here, we combine a frequency-dependent evolutionary matrix game of cooperation with an experiment that makes use of mutant peas which cannot associate with mycorrhizae. Using a model, and an experiment designed to test the model, we ask: (1) what conditions are theoretically necessary for the mycorrhizal association to be an evolutionarily stable strategy (ESS)? We then test the model using a unique mutant of pea (*Pisum sativum* L. R25 (*Pssym8*)) which has been shown to be incapable of forming arbuscular mycorrhizal associations (Balaji *et al*. 1994; Guinel & Geil 2002). Using the experiment, we ask: (2) how does mycorrhizal association across frequency-dependent competitive contexts change reproductive and vegetative yields?; (3) how does the competitive context change the mycorrhizal colonization of wildtype pea roots?; and (4) how does the mycorrhizal association across competitive contexts alter N uptake? Finally, (5) we combine the model and experiment to estimate the costs and benefits of gathering nutrients with and without the mycorrhizal association by solving model equations. Combined, we illustrate how the benefits of mutualism in this case were indeed frequency-dependent; moreover, without the unique mutant, we would have concluded that there were no benefits to this mutualism.

## METHODS

### Evolutionary game theoretic Model

Evolutionary game theory is useful for examining frequency-dependent interactions. Following the logic of classic evolutionary games (e.g. the prisoners’ dilemma Axelrod & Hamilton 1981; Poundstone 1992), we derived a simple matrix game with benefits and costs that allowed us to conceptualize the competition between the R25 mutant and the wildtype (WT) cv. Sparkle based only on their mutualistic associations.

Let *N*_*p*_ be the nutrients that are plant-available, and let *N*_*m*_ be the nutrients that are fungus-available and traded to the plant. We assume *N*_*m*_ is derived from a separate pool of organic nutrients that the plants cannot access without the aid of the mycorrhizal fungus (e.g. Hodge 2001; Leigh, Hodge & Fitter 2009). Similarly, let *c*_*p*_ be the cost associated with nutrient uptake by the plant (e.g. root tissue production, ATP synthesis) and let *c*_*m*_ be the cost to plants of obtaining nutrients via mycorrhizal association (e.g. carbohydrate or lipid trade; (Rich *et al*. 2017)). Since our goal was to consider competition within our own pot experiment, the model considers competition between two plants with a fixed pool of resources, but one could extend this to any number of plants by replacing 2 in any model equations with *x*, where *x* is the number of individuals interacting.

The two pea genotypes (R25 and WT) create four possible combinations of pair-wise competition. First, when two R25 mutants compete, they only have access to plant-available nutrients and we assume that on average each individual plant would access half of those nutrients. R25 plants only pay the cost of obtaining nutrients that are plant-available (Fig 1a). Second, when R25 competes with WT, the outcome for the R25 plant is the same: they compete and obtain only half the available nutrients, and pay the cost of getting those nutrients themselves (Fig 1a). However, when WT competes with R25, it has access to two nutrient pools. WT competes for the plant-available pool of nutrients with R25 obtaining only half on average and paying the cost of obtaining those nutrients, but WT also has access to 100% of the potentially tradeable pool of nutrients accessible by the fungus, and incurs the cost associated with trading for those nutrients (Fig 1a). Finally, when two WT plants compete, they share both nutrient pools on average, and pay both costs.

**FIG 1:**
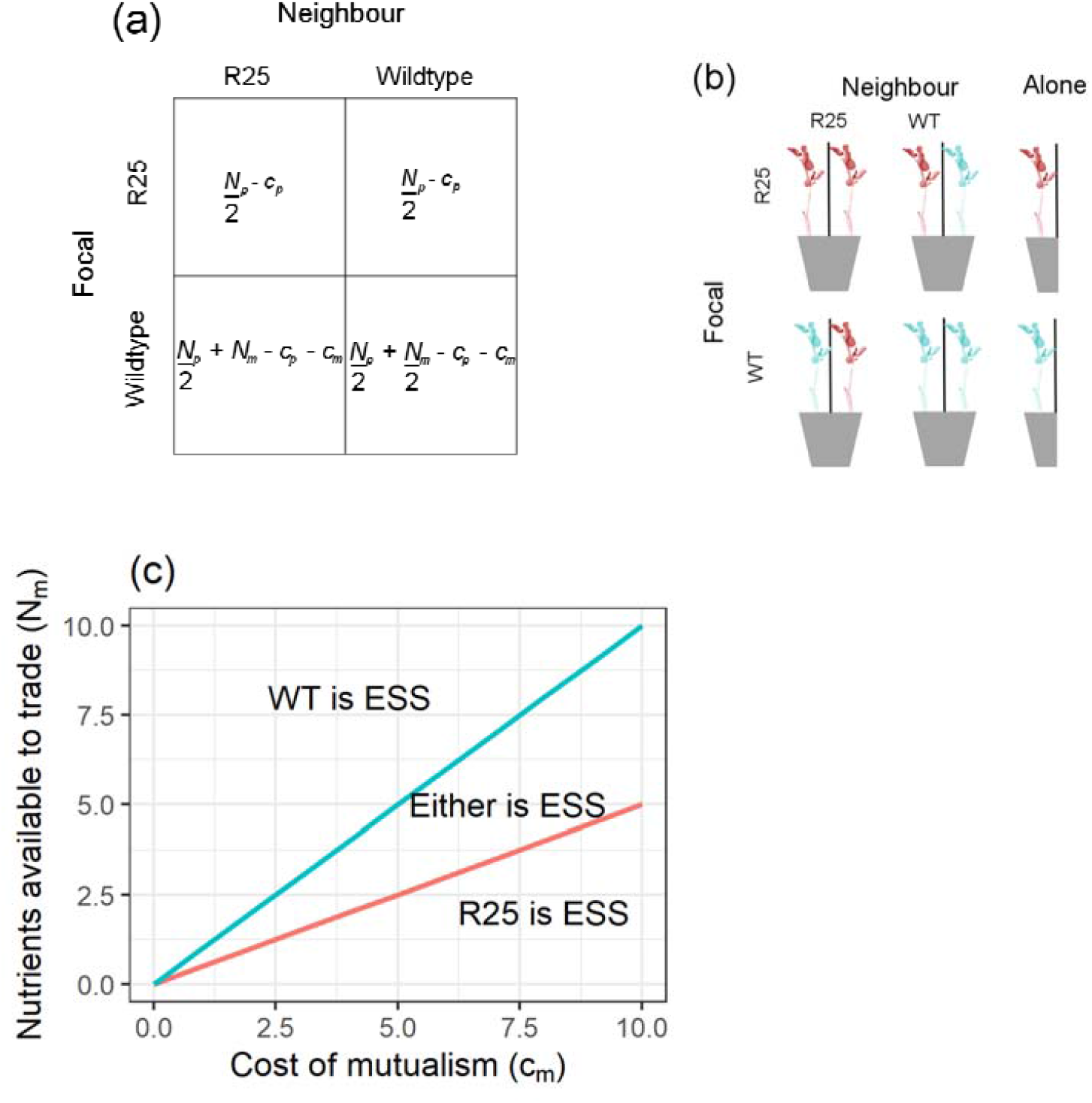
Basic frequency-dependent matrix game for genotype competition (a) and schematic of experimental design (b). Grey trapezoids represent pots, and black vertical lines represent corregated plastic screens erected to prevent above ground interaction. R25 mutant plants are coloured red, and WT plants are coloured blue. There were four competition treatments representing all possible pairs of mutant and WT as either focal and neighbour as captured by the matrix game. Additionally, each plant was grown alone in pots of half the size as a per-plant nutrient control. Thus, there were six treatments in total. The ESS solutions to the matrix game for all possible values of parameter space is shown (c). The blue line represents *N*_*m*_ = *c*_*m*_, and the red line represents 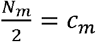. When the costs of mutualism (*c*_*m*_) are greater than half the benefits 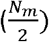, plants should never engage in mutualism (R25 is ESS). When the costs of mutualism (*c*_*m*_) are less than the maximum benefits of mutualism (*N*_*m*_), then plants should always engage in mutualism (WT is ESS). In the limited set of nutrient conditions where *N*_*m*_ > *c*_*m*_, but 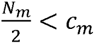, either strategy can be ESS depending on the history of mutations, but they cannot coexist. Instead, a priority effect will favour whichever genotype evolved first within the region and this is defined by *N*_*m*_ > *c*_*m*_, but 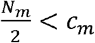.

These interactions described above can be summarised in a two by two payoff matrix (Fig 1a) and by the following equations:

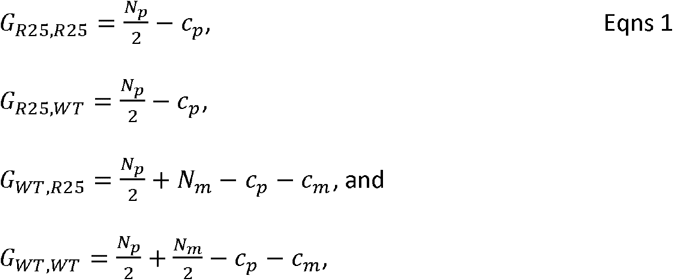

Where *G*_*i,j*_ is individual fitness of focal plant i competing against plant j in each of the four possible pairwise interactions between R25 and WT.

### Evolutionary stable strategy definition

In a matrix game, an ESS is identical to a Nash equilibrium (Maynard Smith & Price 1973; Apaloo *et al*. 2014). An ESS is a strategy (or strategies) which once adopted by members of a population leads to that population not being invaded by any alternative strategy (or strategies). Mathematically, if we imagine that fitness is a function of strategies such that G(v, u) is the fitness payoff of a focal player using strategy ‘*v*’ against a neighbour using strategy ‘*u*’ such that *v* + *u*, then v is a pure ESS if and only if: *G*(*v, v*) > *G*(*u, v*) and *G*(*v, u*) > *G*(*u, u*). Alternatively, *u* is a pure ESS when the inequalities are reversed. Importantly, under this definition, mixed ESS solutions are possible where the two strategies may coexist (*G*(*u, v*) > *G*(*v, v*) and *G*(*v, u*) > *G*(*u, u*)). In addition, priority effects are also possible where both strategies are ESS, but only one can occur at a time depending on the history of mutation (*G*(*v, v*) > *G*(*u, v*) and *G*(*u, u*) > *G*(*v, u*)). Importantly, the ESS can theoretically exist whether or not the exact equilibrium is ever achieved in nature (Maynard Smith 1982).

### Plant material

To examine the effects of mycorrhizal association on competition in a frequency-dependent setting, we took advantage of the pea mutant R25 which has been characterized as a non-mycorrhizal mutant. The R25 mutant was obtained from a background of the cultivar ‘Sparkle’ and was identified from a screen of gamma-radiated seeds (Markwei & LaRue 1992). Remarkably, the non-mycorrhizal phenotype is controlled by a single locus (*sym8*) that exhibits simple Mendelian dominance (Balaji *et al*. 1994). This means that the mycorrhizal and non-mycorrhizal plants, which we will use in our comparison, have a constant genetic background save for a single allele difference. This is exactly the type of evolutionary situation most models envision, where a rare mutant emerges and potentially invades a resident population. This allows us to test evolutionary theory in a novel way.

### Fungal material

The AM fungal strain used in this study was *Rhizophagus irregularis* ((Blaszk., Wubet, Renker & Buscot) C. Walker & Schuessler 2010 as [‘irregulare’]) cv. DAOM 197918 originally obtained from the Agriculture and Agri-Food Canada Glomeromycota in vitro collection (AAFC, Ottawa, ON, Canada) and propagated by FCG using leek (*Allium ampeloprasum* (L.) cv. *muhlenbergii*) as a host. At maturity, leek roots were assayed for mycorrhizal structures and, if present, the leek soil was stored dry at 4°C.

### Experimental treatments

There were four competition treatments that match the four cells of the matrix game (Fig 1a). Treatments included: (*i*) WT versus WT; (*ii*) WT versus R25; (*iii*) R25 versus WT and; (iv) R25 versus R25 (Fig 1b). In each competition treatment, one of the two plants was randomly designated *a priori* as ‘focal’, and the other as ‘neighbour’. With this design, neighbour plants were only present to impose competition, and only the focal plants would eventually be harvested. Thus, even though the R25 versus WT and WT versus R25 treatments were essentially identical, we planted both and measured them independently to avoid pseudoreplication. In addition to the competition treatments, there were also two controls where plants of each genotype were grown alone (Fig 1b).

There is some debate in the literature on the appropriate control in competition experiments. Should one control for the nutrient environment (i.e. total nutrients provided per plant, and soil nutrient concentration; (e.g. Gersani *et al*. 2001; McNickle & Brown 2014; Chen *et al*. 2015a)), or should one control for volume but not for the nutrient environment (e.g. Hess & De Kroon 2007; Chen *et al*. 2015a)? Unfortunately, it is difficult to control both nutrients and pot volume without confounding one of them with neighbour addition (McNickle *in press*), and thus one assumption of our design was that both nutrient availability and soil nutrient concentration are more important for belowground plant-plant competition than pot volume. Thus, in total there were four competition treatments (all possible pairs of R25 and WT), and two no-competition controls (two genotypes grown alone) replicated 15 times for a total of six treatments and 90 pots (Fig 1b).

### Greenhouse conditions

All plants were grown in 15-cm diameter, 15-cm high, standard plastic pots in the Wilfrid Laurier University greenhouse in Waterloo, Ontario, Canada (43°28’28.1”N, 80°31’15.2”W) from June 1 to July 29, 2015. Artificial light was not used yielding approximately 15h:9h light:dark at this latitude and time of year. Temperature was maintained at approximately 25°C. The model assumed that all interactions occurred belowground, thus screens of corrugated white plastic were erected in the middle of each pot so that plants could not interact above-ground (Fig 1b). Soil was a 1:1 mixture of peat moss (Greenworld Garden Products, Pointe-Sapin, New Brunswick, Canada), and calcined clay gravel (Turface® MVP, PROFILE Products LLC, Buffalo Grove, Illinois, USA). In addition, we added to the soil, at a rate of 0.01% by volume, ground WT pea shoots which had been labelled beforehand with elevated ^15^N by growing them with labelled ammonium nitrate. Our rationale for this was that organic sources of nitrogen are known to be more readily available to the fungus than to the plant (Hodge 2001; Leigh, Hodge & Fitter 2009; Hodge & Fitter 2010), and the isotopic signature of the plants would let us determine nitrogen trade. This entire soil mixture was then autoclaved at 120°C for 20 minutes in 10L batches.

Seeds of each pea genotype were surface-sterilised in 0.4% sodium hypochlorite for ten minutes, rinsed three times with deionized water, and then soaked in deionized water for 18 hours prior to planting. Only those seeds that had swollen and imbibed water were planted to maximize germination. To inoculate pots, the sterile soil mixture described above was mixed with soil from the leek trap cultures at a ratio of 1:9. Since R25 cannot form associations with mycorrhizae, all soil in the experiment was inoculated with live culture as a control, even soil on which the non-mycorrhizal R25 would eventually grow.

Nutrients were added as a mineral nutrient solution (Miracle Grow® all-purpose water soluble plant food, Scotts Canada Ltd, Mississauga, Ontario, Canada). Plants were fertigated with 200mL of 0.5g/L nutrient solution every 7 days on Monday afternoons and then irrigated with 200mL of water every 7 days on Friday mornings. Fertigation began when the plants were 14 days old; so, for the first 14 days plants only received water on Mondays and Fridays. Each pot in the entire experiment was placed in its own individual circular tray. The trays collected any water or nutrients that drained through the pot making each pot its own closed nutrient system, and where water was only lost through evaporation or transpiration. Pots were arranged in a randomized block design to control for potential microclimate effects within the greenhouse. In addition, because of the screens erected to block competition for light, pots were turned one quarter turn every Monday and Friday to minimize the potential effects of any shading from the screens. The appropriate nutrient concentration used was determined through a pilot experiment (Supplementary information).

### Harvest

All plants were grown for 60 days until senescence began, and were then harvested. At harvest, above ground material was sorted into shoots and fruits. These samples were dried at 60°C and weighed. Below ground, fourteen 3cm-long root fragments were randomly sampled from each focal plant in each pot; they were stained to assess mycorrhizal colonization as described below. In the competition treatments, we could work down from the stems of each plant to ensure that we collected only root fragments that belonged to focal plants. These root fragments were air-dried until staining occurred. The remainder of the roots were washed on a 2mm sieve, dried and weighed. For total root biomass, we attempted separation but were unable to separate the root systems of the two plants.

### Estimation of mycorrhizal colonisation

Dried root fragments were rehydrated in deionized water for 25 minutes in 1.5mL centrifuge tubes before staining (Vierheilig *et al*. 1998). Briefly, root fragments were cleared in 10% KOH in two steps (one at 95°C for 10 minutes, and the other at 95°C for 5 minutes). Cleared root fragments were rinsed twice with 5% acetic acid and stained with a 1:20 (v:v) mixture of black Indian ink (Scheaffer Pen and Art Supply Co., Providence, Rhode Island, USA) and 5% acetic acid at 95°C for five minutes. Fragments were de-stained with 5% acetic acid for 18 hours, and vacuum-infiltrated in glycerol (30% and 60%, 20 minutes each); on each slide, 7 root fragments were mounted in 60% glycerol. In total, we collected 14 fragments per plant and thus there were two slides per plant for a total of 240 slides.

Mycorrhizal colonisation was scored according to the magnified intersections method described in McGonigle *et al*. (1990). All intersections between the root and the eyepiece cross hair were examined for colonisation, and the depth of view was adjusted to examine the entire root. At each intersect we counted the number of (a) arbuscules; (b) vesicles; and (c) intra-radicular hyphae; we also noted any absence of mycorrhizal structure. These data allowed us to calculate the proportion of root length colonized by each of the three mycorrhizal structures.

### Stable isotope analysis

Shoot material (leaves and stems) of focal plants were ground to a fine powder using a bead mill (Mixer Mill MM400, Retsch GmBH, Haan, Germany). Approximately 3mg of ground tissue was weighed into tin capsules (Part number 041061, Costech Analytical Technologies Inc, Valencia, CA, USA). These samples were then processed by the University of California Davis Stable Isotope facility using a PDZ Europa ANCA-GSL elemental analyzer interfaced to a PDZ Europa 20-20 isotope ratio mass spectrometer (Sercon Ltd., Cheshire, UK). Importantly, and perhaps counter-intuitively, due to fractionation of ^15^N, lower levels of enrichment indicate more trade.

### Data analysis

In general, all data were analysed using general linear mixed effects models (GLMM) in R (v3.3.2) using the lme4() and lmerTest() libraries, a type III sum of squares, and the Satterthwaite approximation for denominator degrees of freedom when needed (Bates 2007; R-Development-Core-Team 2009). To find adequately fitting models, we first tried to adjust to alternative probability distributions, and failing that, we transformed data if necessary. A separate GLMM was fit for: i) shoot, fruit, and root mass; ii) percent colonisation of intraradicular hyphae, arbuscules and vesicles, and; iii) ^15^N enrichment, and N concentration as detailed below for a total of eight analyses.

First, analysis of biomass data included the full factorial combination of treatment (mixed genotypes grown together, same genotypes grown together, or plants alone) by genotype (WT or R25 mutant) with block as a random intercept. Shoot and root biomass was analysed using a Gaussian error distribution. Fruit biomass required log(*x* + *c*) transformation to achieve an adequate model fit using a Gaussian error distribution, where c was half the smallest fruit mass measured. Significance was assessed using an F ratio test and a type III sum of squares.

Second, mycorrhizal colonisation was analysed with only treatment (mixed genotypes grown together, same genotypes grown together, or plants alone) and microscope slide nested inside block as a random intercept since two slides were separately scored per individual. Since these were count data which were then used to calculate a proportion of root length colonized, we analysed them as a binomial GLMM where the response variable was the proportion of root length colonised, with the logistic regression weighted by the total number of observations (Harrison 2015). Significance was assessed with a type III Wald’s Chi, which is appropriate for binomial mixed models (Bolker *et al*. 2009).

Finally, the nitrogen data were analysed in a manner similar to the biomass data with the full factorial combination of treatment (mixed genotypes grown together, same genotypes grown together, or plants alone) by genotype (WT or R25 mutant) with block as a random intercept. Although N concentration and ^15^N enrichment were both proportion data, unlike the mycorrhizal data they were not based on counts, making a binomial distribution inappropriate. We thus arcsine square root transformed both N variables for continuity to achieve adequate model fits with a Gaussian distribution.

## RESULTS

### Model

From the ESS definition, and eqns 1, then we find that R25 is the ESS when:

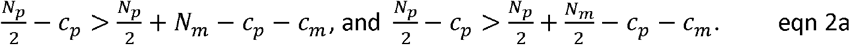

Eqn 2a can be simplified to:

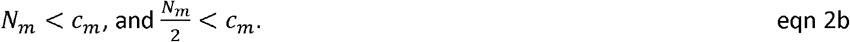

Similarly, WT is ESS when the opposite conditions to eqn 2a are met,

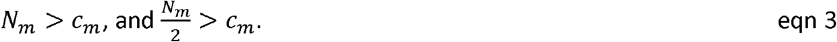

Interestingly, there is only one more possible solution to this matrix game, which can be interpreted as alternative stable states, where both strategies are ESS, but they cannot coexist. Under this solution, priority effects determine which strategy would be found within the population, but only one strategy can exist at a time. This occurs under the limited range of nutrients where each genotype does better in monoculture than in mixture. That is when,

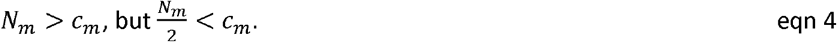

In general, matrix games have a fourth possible solution where the two strategies coexist within a population. However, this solution is not possible in this game because it requires

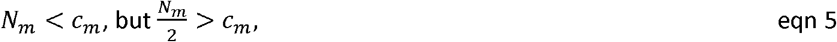

which is logically impossible because 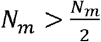

Since *N*_*m*_ and *c*_*m*_ are the only relevant parameters in determining the ESS, we can summarise the conditions in eqns 1-5 graphically in *N*_*m*_ and *c*_*m*_ space showing all possible combinations of nutrient availability and mutualism costs and the resulting ESS solutions (Fig 1c).

### Experiment: plant biomass

Biomass was analysed with the competition treatment (same genotype, mixed genotype or alone) crossed with focal genotype (R25 or WT) in a full factorial design that also included block as a random effect.

Beginning with fruit biomass, which represents life-time reproductive effort (i.e. fitness) in this annual plant, there was a significant interaction between treatment and genotype (Table 1). Post-hoc comparisons revealed that this interaction was driven by the WT genotype grown in competition obtaining significantly more fruit biomass than plants in other treatments (Fig 2a). However, when WT was grown in competition with another WT, the fruit biomass was not statistically different from that of R25 nor was it different from that of either genotype grown alone (Fig 2a). For shoot biomass, only the genotypes were significantly different, and the WT was larger than the mutant in all treatments (Table 1, Fig 2b). For root biomass, there were no significant differences among treatments or genotypes (Table 1, Fig 2c). However, roots of the two competing plants could not be separated; thus in the mixed genotype competition treatments, it is difficult to draw conclusions about root biomass.

**Table 1:** Results of GLMM on plant growth using the lme4() and lmerTest() libraries in R. Denominator degrees of freedom (Den df) were estimated using the Satterthwaite approximation and a Gaussian distribution. The fruit mass data were log(x+c) transformed to achieve model fit, where c was half the smallest measured value. All models included block as a random intercept. Bold with * indicates statistical significance at p<0.05.

| Fruit |  |  |  |  |
| --- | --- | --- | --- | --- |
| Factor | Num df | Den df | F | p |
| Treatment | 2 | 135 | 2.82 | 0.0633 |
| Genotype | 1 | 135 | 9.63 | **0.0023\*** |
| Treatment X Genotype | 2 | 135 | 3.84 | **0.0239\*** |
| Shoot |  |  |  |  |
| Factor | Num df | Den df | F | p |
| Treatment | 2 | 143.0 | 0.05 | 0.9555 |
| Genotype | 1 | 143.0 | 6.95 | **0.0093\*** |
| Treatment X Genotype | 2 | 143.0 | 1.22 | 0.2998 |
| Root |  |  |  |  |
| Factor | Num df | Den df | F | p |
| Treatment | 2 | 124.1 | 1.92 | 0.1502 |
| Genotype | 1 | 123.1 | 0.21 | 0.6473 |
| Treatment X Genotype | 2 | 123.2 | 0.06 | 0.9420 |

**FIG 2:**
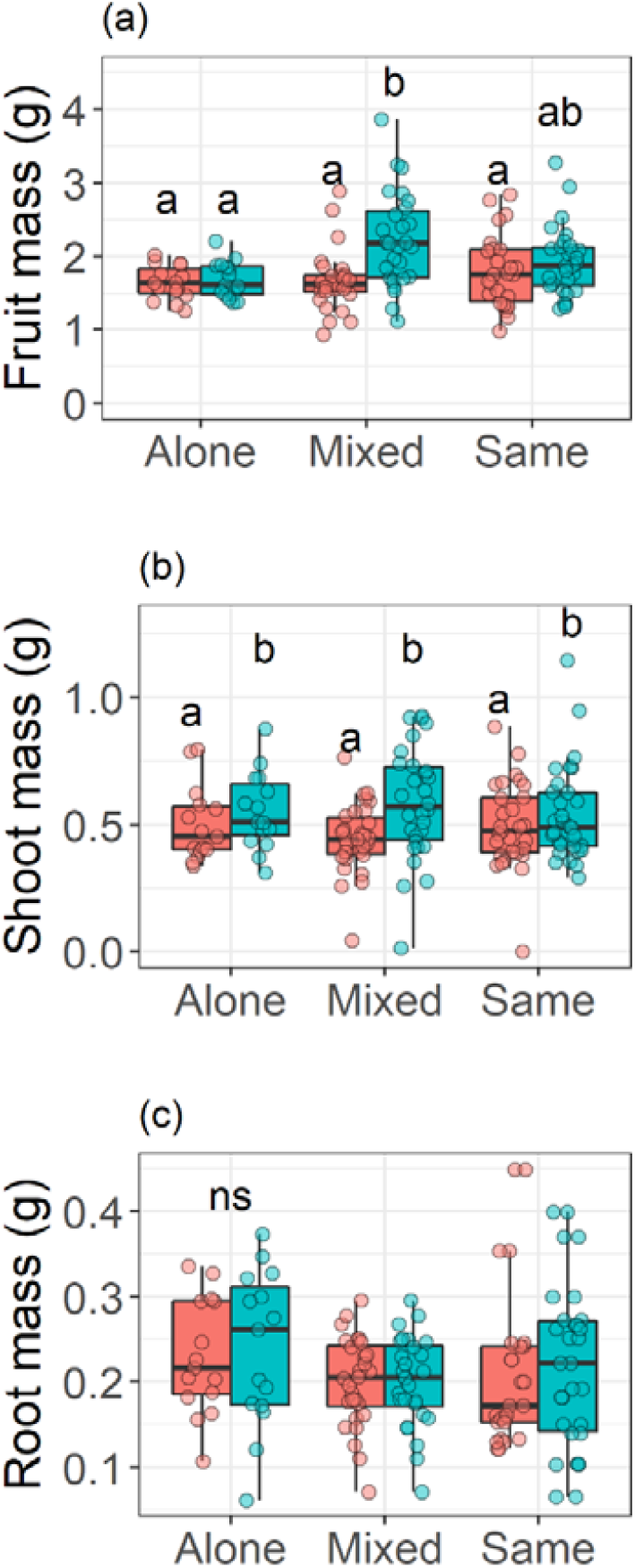
The fruit (a), shoot (b), and root (c) production of plants, with R25 shown in red and WT in blue either grown alone, in mixed genotype competition (i.e. R25 vs WT and WT vs R25), or in same genotype competition (i.e. R25 vs R25, or WT vs WT). Letters indicate significant differences in means from post-hoc comparisons, while ns indicates a lack of statistical significance. The raw data are plotted with a jitter around the boxplots. Note that in (c) we could not separate the roots of plants in competition and so the data shown represent the roots of both plants combined, and the colour represents the genotype of the focal plant even though in mixed treatments the roots of the neighbour were also weighed.

### Experiment: mycorrhizal colonisation

Since we confirmed that the R25 mutant is physiologically incapable of forming an association with arbuscular mycorrhizal fungi because of a mutation in the *sym8* locus (Fig S3; Balaji *et al*. (1994)), the analysis only included the colonisation of the WT roots across the competition treatments (alone, mixed genotype, or same genotype) with block as a random effect. There were no significant differences in the proportion of root length colonised by either the intraradicular hyphae (χ^2^=3.04, df=2, p=0.2181, Fig 3a) or the arbuscules (χ^2^=2.02, df=2 p=0.3634, Fig 3b). However, there was a significant difference in vesicles among the treatments (χ^2^=328824.0, df=2, p<0.0001, Fig 3c). All three treatments were different where plants grown alone had the highest number of vesicles, followed by plants grown in mixed genotype competition, and finally by WT competing with WT plants.

**FIG 3:**
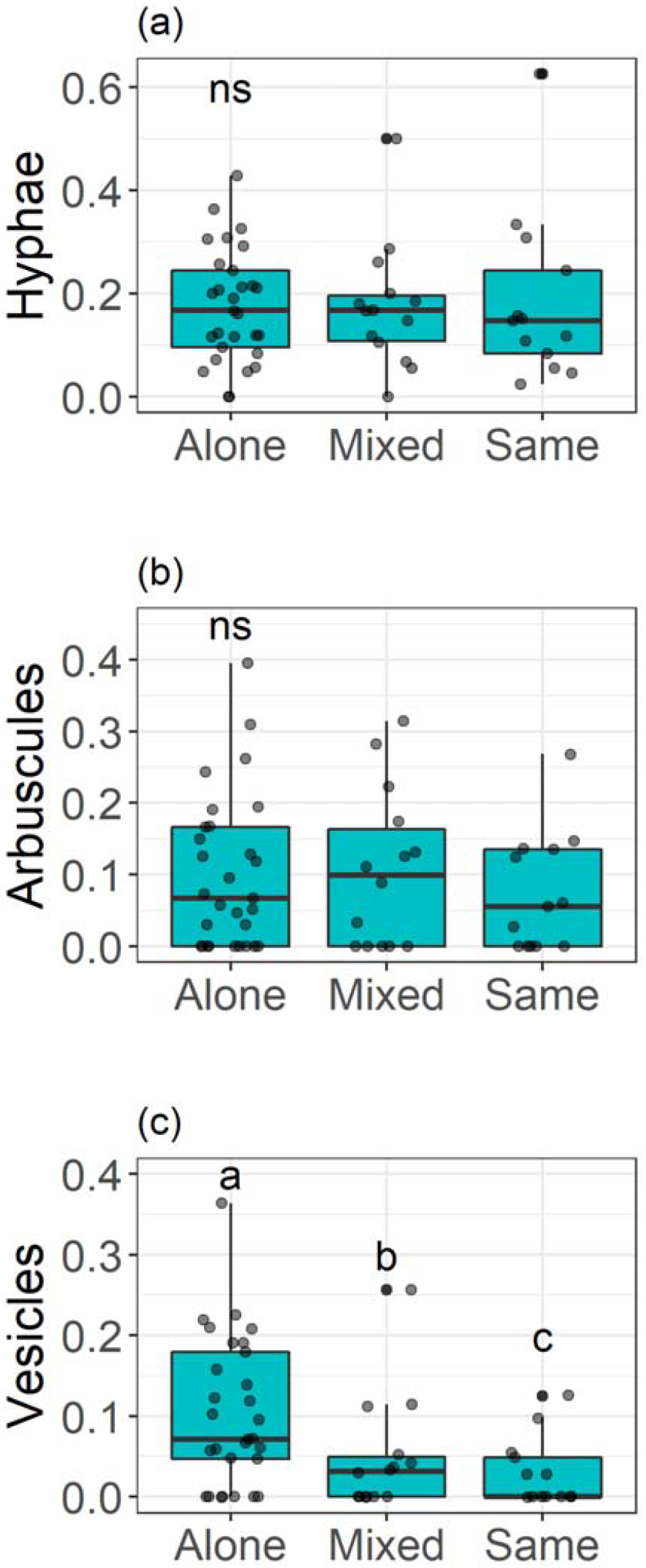
The colonisation of roots as a proportion of length by the mycorrhizal structures of (a) intraradicular hyphae, (b) arbuscules and (c) vesicles. Note R25 is incapable of forming mycorrhizal associations (Fig S3), and so the data shown are only for WT plants. The raw data are plotted with a jitter around the boxplots. Letters represent significant differences among the treatments on the x-axis and ns indicates no statistical significance.

### Experiment: nitrogen trade

Both ^15^N enrichment and shoot N concentration were analysed with the competition treatment (same genotype, mixed genotype or alone) crossed with genotype (R25 or WT) in a full factorial design with block as a random effect. For ^15^N enrichment, the main effects of treatment and genotype were statistically significant, but their interaction was not (Table 2). Thus, we plotted these data for each treatment and genotype (Fig 4a), as well as for each genotype with all treatments combined (Fig 4b) to show most clearly the main effects in the absence of an interaction. Post-hoc tests revealed that plants that experienced competition of any type were more enriched in ^15^N than plants grown alone (Fig 4a), while the R25 mutant was more enriched than the WT (Fig 4b) as expected due to fractionation during mycorrhizal trade.

**Table 2:** Results of GLMM on %^15^N enrichment and nitrogen concentration (% mass/mass) of plant shoot material using the lme4() and lmerTest() libraries in R. Denominator degrees of freedom (Den df) were estimated using the Satterthwaite approximation. All models included block as a random effect.

| % <sup>15</sup>N enrichment |  |  |  |  |
| --- | --- | --- | --- | --- |
| Factor | Num df | Den df | F | p |
| Treatment | 2 | 65.7 | 22.28 | <0.0001 |
| Genotype | 1 | 66.0 | 9.73 | 0.0027 |
| Treatment X Genotype | 2 | 65.9 | 0.08 | 0.9198 |
| N Concentration |  |  |  |  |
| Factor | Num df | Den df | F | p |
| Treatment | 2 | 65.7 | 1.71 | 0.1886 |
| Genotype | 1 | 65.8 | 4.34 | 0.0403 |
| Treatment X Genotype | 2 | 65.7 | 1.68 | 0.1951 |

**FIG 4:**
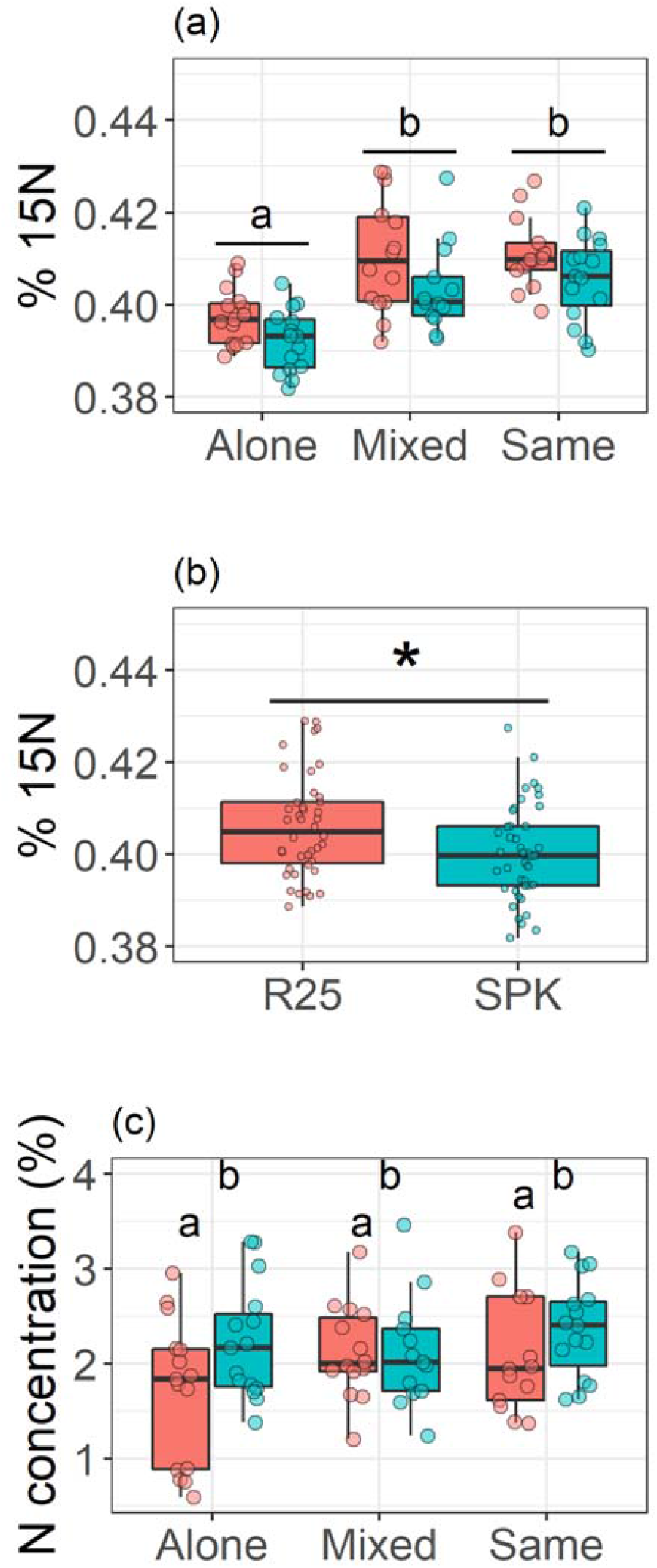
Shoot stable isotope enrichment across both treatments and genotypes (a), as well as just among genotypes (b). The total N concentration of shoots is also shown in percent (mass/mass) (c). WT is shown in blue and the mutant R25 in red. In each panel, the raw data are plotted with a jitter around the boxplots. Letters or * represent significant differences among the treatments on the x-axis. In panel (b), the genotypes are highlighted because the interaction between treatment and genotype was not significant for %15N enrichment (Table 2), and thus pooled genotypes 15N enrichment was used to parameterize the model.

For total N concentration, only the genotype was statistically significant, and the WT had higher N content than the non-mycorrhizal mutant (Table 2, Fig 4c). This indicates that mycorrhizal trade led to fitness benefits directly through increased N.

### Combining model and experiment

From our harvest data, we can use the fruit mass of these annual plants as an estimate of life-time fitness payoffs (*G*) to solve for the parameter values required to determine the ESS. To do this, we combined means according to the GLMM and post-hoc analyses (Table 1, Fig 2a) to obtain the payoff matrix shown in Table 3.

**Table 3:** Parameterized values for *G* in each interaction estimated via mean lifetime reproductive yield of all plants.

|  |  | Neighbour |  |
| --- | --- | --- | --- |
|  |  | R25 | WT |
| Focal | R25 | 1.73 (95% CI: 1.61 – 1.85) | 1.73 (95% CI: 1.61 – 1.85) |
|  | WT | 2.23 (95% CI: 1.99 – 2.47) | 1.93 (95% CI: 1.76 - 2.10) |

To estimate *N*_*p*_ we begin with the fact that we know that the R25 mutant obtained all of its N through direct uptake as *N*_*p*_, while the WT obtained some N itself as *N*_*p*_ and some through trade as *N*_*m*_ (Fig S3, Fig3b). The pea genotypes significantly differed in the N concentration of their tissues with the WT having significantly higher N concentrations than the mutant by an average of 2.7%. We assume that the other 97.3% of N in the WT tissues must have come as *N*_*p*_. Thus, if we assume that WT plants have the same *N*_*p*_ as R25 plants, then we know that:

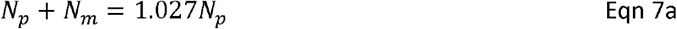

And therefore that

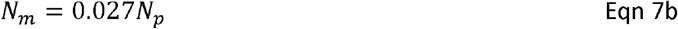

Substituting mean fitness as *G* and *N*_*m*_ = 0.027*N*_*p*_ into eqns 1 gives us the following three equations with three unknowns that can be rearranged to find:

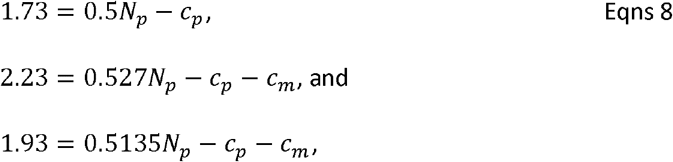

The solutions to this set of equations are *N*_*p*_ = 22.2 (and therefore, *N*_*m*_ = 0.6 via eqn 7b), *c*_*p*_ = 9.4, and *cm* = 0.1. From eqns 2 and 3, the mycorrhizal association is expected to evolve when, 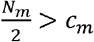 and since 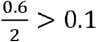, then mycorrhizal mutualism is an ESS within our model and experimental system. It might seem trivial for our model to return what we already knew: that the wildtype was the ESS, and the mutant artificially created by gamma radiation was not ESS. However, the value of a validated model which we have failed to reject is that it can be exported to other systems to make novel predictions.

## DISCUSSION

Evolutionary models suggest that mutualism evolved because it conferred an advantage to mutualistic individuals over non-mutualistic individuals in a population, and thus it became the ESS (Axelrod & Hamilton 1981). However, such models are not always easy to empirically validate and test. The game theoretic model we presented here which was specific to the R25 mutant pea system supported this classical finding. Previous studies of plant-plant competition with and without mycorrhizal associations have generally found that either competing plants perform worse in the presence of the fungi (Fitter 1977; Hodge 2003), or that there is no difference among competing plants with or without mycorrhizal partners (Allen & Allen 1984; Moora & Zobel 1996). Indeed, if we had only considered R25 vs R25 and WT vs WT treatments in our study, though the shoots of WT were larger than R25, we would have also concluded that there was no fitness advantage to the mycorrhizal association (Fig 2a). However, the use of the R25 mutant allowed us to examine the full frequency-dependent context, and it revealed that the mycorrhizal WT had an advantage in terms of reproductive output only when it competes with a non-mycorrhizal R25 mutant. When this result is placed within the fitness payoff matrix of the evolutionary game, it reveals that mutualism is an ESS even though there was no difference in either pure population.

At first, the lack of difference in fitness within the R25 vs R25 treatment and WT vs WT treatments might seem like evolution should be indifferent to whether plants form mycorrhizal associations, but the key to frequency-dependence is its context-dependence. Using the notation from our definition of ESS, we could write this result as *G*_*R25,R25*_ = *G*_*WT,WT*_, but note that this comparison is not relevant to the definition of the ESS that requires either *G*_*WT, WT*_ > *G*_*R25,WT*_ and *G*_*WT,R25*_ > *G*_*R25,R25*_, or the opposite to achieve one of the two possible pure ESSs. That is, the relative fitness of either pure population is entirely irrelevant to the question of what trait should be favoured by natural selection. Since the ESS is about rare mutants invading pure populations, or pure populations resisting invasion (Maynard Smith & Price 1973; Maynard Smith 1982), we cannot over emphasize how important the mixed genotype (i.e. R25 vs WT and WT vs R25) treatments are to understanding the frequency-dependent benefits of mutualism.

When combined, the model and the experiment allowed us to generate estimates of the costs and benefits of mycorrhizal association. These calculations express both costs and benefits in units of fitness under the assumption that N uptake and fitness are correlated (Fig S1c). We estimated all the model parameters, but the most interesting is to compare 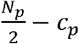 and 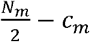, which represent the net-benefit obtained by the plant’s own foraging behaviour versus the net benefit in increments of fitness obtained from association with mycorrhizal fungi, respectively. Since 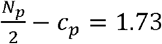, and 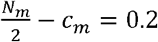, this means that mycorrhizal plants in this experiment gained approximately 11% of their reproductive output as a direct benefit of the mutualistic association with mycorrhizal fungi. In the WT vs R25 treatments, this 11% benefit manifested as a significant increase in reproductive output (Fig 2a). However, in the WT vs WT treatment, both plants had this benefit, and so it was not apparent (Fig 2a).

A key caveat to these conclusions is that we assume that *N*_*p*_ remains unchanged in both R25 and WT genotypes. This requires that the plant does not adjust its own foraging activities when mycorrhizae are present. We feel that this is valid because the two genotypes differed only by one allele at a single locus. Another caveat is that interactions between two plants are far from a population-level context. Indeed, competition among two plants is the absolute minimum design to examine such questions, but a range of population densities would allow a stronger test (Hart, Freckleton & Levine 2018).

So far, we have only discussed the plants, but there are two partners in this association and it is worth considering the success of the fungal partner. Arbuscule and intraradicular hyphae colonisation did not vary among treatments, but we found a significant effect of our competition treatments on the colonisation of vesicles inside pea roots such that the most vesicles were contained in plants grown alone, fewer in the mixed genotype competition pots, and fewer still in the same genotype competition pots (Fig 3c). Vesicles are storage organs of the fungi that accumulate lipids and can also become propagules upon root death (Biermann & Linderman 1983). Thus, one interpretation of our data is that potential fungal propagules (an admittedly weak surrogate of fungal fitness) also declined along a gradient of increasing competition intensity for the plants. This might suggest that there was increased fungal competition when the fungus was connected to more than one host. This could be similar to a tragedy of the commons game among plants, where plants over-proliferate tissues to maximize their competitive ability (Gersani *et al*. 2001; Mcnickle & Dybzinski 2013). This over-proliferation occurs in roots, leaves and stems and, while ESS maximizes competitive ability, it tends to reduce reproductive output (Dybzinski *et al*. 2011; Mcnickle *et al*. 2016). Our results may suggest that the fungal partner, acting as an extension of the root system, is also increasing its number, and length, of hyphae to maximize its competitive ability. This allocation of resources towards growth of the extraradicular hyphae in the soil could be at the expense of vesicle differentiation, as is indicated by the reduced number of vesicles (Fig 3c). The evidence for this is weak, and we have no data about the fungus behaviour outside of plant roots, but we suggest this is an interesting hypothesis that requires future attention.

Related to this possible tragedy of the commons response in the fungi, we highlight that no over-proliferation leading to a tragedy of the commons was observed for the roots. This seems to be a consistent finding for pea (Meier *et al*. 2013; Chen *et al*. 2015b) (McNickle in press, but see (O’Brien, Gersani & Brown 2005)).

### Conclusion

Here we combined an evolutionary game theoretic model and an experiment with loss of function pea mutants that could not form mycorrhizal associations to show that the benefits of mycorrhizal cooperation are frequency-dependent. We used a mutant pea genotype that differs by one allele from the WT to examine the full factorial evolutionary context contained in models (Fig 1). First, we showed that mycorrhizal association would be the ESS when half the potential fitness benefits are greater than the costs of the association (Fig 1c). Second, we showed that the reproductive output of mycorrhizal plants was only significantly greater than that of non-mycorrhizal plants in the frequency-dependent context of WT versus mutant (Fig 2a). Third, the only fungal structures that varied across competition treatments were the vesicle storage organs, the number of which declined with increasing competition intensity of plants (Fig 3c). Finally, we combined the experiment and the model to calculate the costs and benefits of plant nutrient capture and plant trade with the mycorrhizal fungus; we estimated that 11% of mycorrhizal plant fitness was directly attributable to the mutualism. Importantly, without the R25 mutant that is incapable of forming mycorrhizal associations, we would have concluded that mycorrhizae conferred no advantage. This result highlights the importance of frequency-dependent selection in ecology and evolution. We propose that what has been commonly called the mutualism-parasitism continuum of mycorrhizal associations (Johnson, Graham & Smith 1997) might be better described as a more beneficial – less beneficial continuum.

## ACKNOWLEDGEMENTS

Funding came from the Natural Sciences and Engineering Research Council of Canada for a Banting postdoctoral fellowship to GGM, and Discovery funding to JLB; from Wilfrid Laurier University to JLB and FCG; from the Ontario Graduate Scholarship programme to AES; and from Purdue University and USDA-NIFA Hatch funds, Grant/Award Number: 1010722 to GGM.

