## SUPPLEMENTARY INFORMATION for "The Benefits of Mycorrhizae are Frequency-Dependent: A Case Study With a Non-Mycorrhizal Mutant of *Pisum Sativum*"

A separate experiment was performed to examine the effects of different fertilizer levels on pea growth and mycorrhizal association. This was done for two reasons. First, competition below ground is expected to be the most intense when mineral nutrients are more limiting than carbon (Chapin, Vitousek & Vancleve 1986; Mcnickle *et al.* 2016). Second, mycorrhizal associations are known to be sensitive to nutrient availability. Specifically, mycorrhizal associations tend to decline as nutrient levels increase (Johnson, Graham & Smith 1997).

**METHODS**:

All conditions were identical to the main experiment described in the main text, and the experiment was run concurrent with the main experiment. Pea plants were grown in 15cm diameter, 15 cm high standard plastic pots in the Wilfrid Laurier University greenhouse in Waterloo, Ontario (43°28'28.1"N 80°31'15.2"W) from June 1 to July 29, 2015. Artificial light was not used so the light level and photoperiod was determined by sunrise and sunset (approximately 15:9 light:dark at this latitude and time of year) and temperature was maintained at approximately 25°C. Soil was a 1:1 mixture of peat moss (Greenworld Garden Products, Pointe Sapin, New Brunswick), and calcined clay gravel (Turface® MVP, PROFILE Products LLC, Buffalo Grove, Illinois, USA). This soil mixture was then autoclaved at 120°C for 20 minutes in small batches to sterilize. Mycorrhizal inoculum of *Rhizophagus irregularis* (N.C.Schenck & G.S.Sm.) was obtained from trap cultures in association with leek (*Allium ampeloprasum* L., var. Musselburgh) grown in the same soil described above. The 1:1 mixture of gravel and peat was mixed with soil from the trap cultures at a ratio of 1:9. Seeds were surface sterilised in 0.4% sodium hypochlorite for ten minutes, rinsed three times in sterile deionized water, and then soaked in water for 18 hours prior to planting. Only swollen seeds were planted to ensure germination. There was only one plant per pot.

In this experiment, the only treatment was nutrient addition. Nutrients were added at five different concentrations along a log scale. The nutrient concentrations were: 0.25, 0.5, 1, 2, and 4 g/L of a mineral nutrient solution (Miracle Grow® all-purpose water soluble plant food, Scotts Canada Ltd, Mississauga, Ontario, Canada). Plants were irrigated with 200mL of water every 7 days on Friday afternoons and fertigated with 200mL of nutrient solution every 7 days on Tuesday mornings. Thus, plants received water every 3.5 days. Each pot was placed in its own circular tray. Water could drain from pots into the trays, but not onto the floor so that we could be certain that the nutrients added were available to the plants. We included eight replicates of the two genotypes (Sparkle, and R25) for a total of 80 pots. Pots were arranged in a randomized block design to control for potential microclimate effects within the greenhouse.

Wild type (WT) and mutant (R25) plants were grown for 60 days until senescence began, and were then harvested. At harvest, above ground material was sorted into leaves (including petioles), stems and fruit. Each of these organs was dried at 60°C and weighed. Below ground, fourteen 3cm-long root fragments were randomly sampled from each plant in each pot; these fragments were stained for an assessment of colonization. The remainder of the roots were washed on a 2mm sieve, dried and weighed.

Mycorrhizal colonisation was scored according to the method described in Mcgonigle *et al.* (1990), and exactly as described in the main text.

Data were analyzed using general linear mixed effects models in R (v3.3.2) using the lmer and lmerTest libraries, a type III sum of squares, and the Satterthwaite approximation for denominator degrees of freedom. In each case, the dependent variables were the natural log of nutrient concentration as a continuous variable, crossed with genotype. We performed three main analyses. First, we analysed total root, shoot and fruit biomass; biomass data were log transformed for normality. Second, we analysed the relative allocation to leaves, stems and roots. These data were proportions which can only fall between 0 and 1, and thus we arcsine square root transformed these for continuity. Finally, we also analysed the proportion of root length colonized by fungal arbuscules, vesicles and intra-radicular hyphae. These proportions were also arcsine square root transformed.

**RESULTS:**

*Plant biomass*

As is characteristic and expected of plant growth, shoot biomass (Fig S1a) and root biomass (Fig S1b) both showed a concave down pattern where root biomass reached a maximum at lower nutrient levels than shoot biomass (FIG S1). These different peaks represent the shift from nutrient limitation to carbon limitation. Fruit production reached a peak at an intermediate value of nutrient addition (Fig S1c). The nutrient levels to the left of the peak for root mass represent nutrient limitation (<0.25g/L), the nutrient levels between the peak for root mass and the peak for shoot mass represent carbon limitation (0.5 – 4g/L), and nutrient levels to the right of the peak for root, shoot and fruit biomass would indicate nutrient toxicity (>4g/L; However, toxicity was not observed in this study). In all cases, the effect of nutrient addition on plant growth was statistically significant, but strains only differed in their fruit biomass, with the mutant strain having slightly lower fitness than the wildtype (Table S1, Fig S1).

*Relative allocation*

Similar to biomass, allocation was significantly influenced only by nutrient addition, and did not differ between the two genotypes (Fig S2, Table S2). Allocation shows the typical root-shoot trade-off along a nutrient gradient, where plants invest relatively more into roots than shoots as nutrients become limiting. Interestingly, in pea, this was primarily a leaf – root trade-off (Fig S2). Stem allocation also significantly declined as nutrients increased, but quantitatively this decline was modest (Fig S2).

*Mycorrhizal association*

As expected, the proportion of root length colonised by mycorrhizal structures declined with increasing nutrient availability in the wildtype (Fig S3). As for the R25 pea mutant, consistent with previous literature, little to no mycorrhizal association was detected. A small amount of hyphae was observed in R25 roots (Fig S3c), but importantly these hyphae did not differentiate into arbuscules (Fig S3a), which are the fungal structures essential for the mutualism to function. Unsurprisingly, these two results produced a significant two-way interaction between nutrient addition and genotype (Table S3).

*Implications for main experiment*

The experiment described in the main text exclusively used 0.5g/L as the fertiliser (-0.69 on a log scale). Importantly, this nutrient level is: (i) below the transition from nutrient limitation to carbon limitation that causes plants to shift allocation to shoots (Fig S1a,b, Fig S2); (ii) below any signs of toxicity in terms or reduced reproductive success (Fig S1c,), and; (iii) still resulted in significant colonisation by beneficial mycorrhizal fungi (Fig S3).

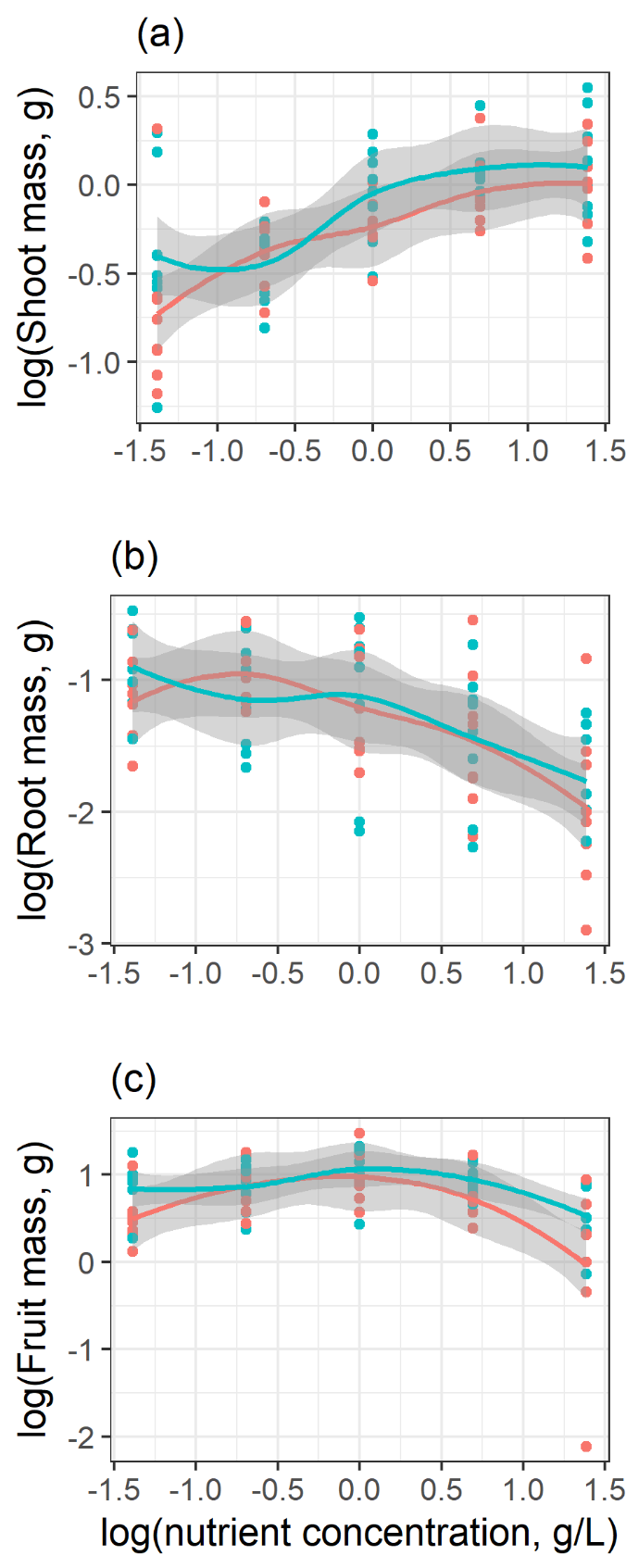

**FIG S1:** Growth and reproduction of mycorrhizal plants along a nutrient gradient. Wild type plants are shown in blue, mutant plants in red. (a) Shoot mass, (b) Root mass and (c) Fruit mass. Lines represent loess regressions, and the grey bands are standard errors.

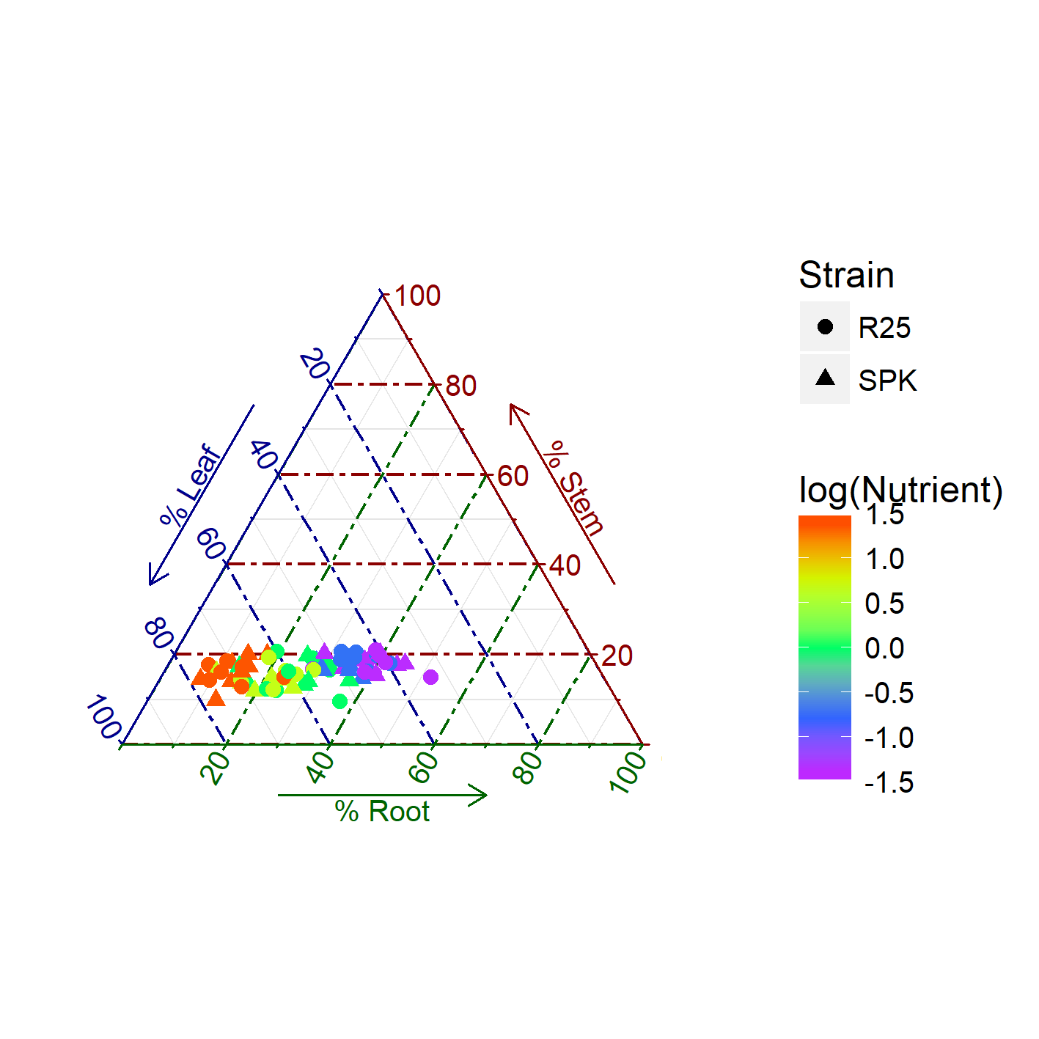

**FIG S2:** Relative allocation to leaves, stems and roots by wildtype (SPK) and mutant (R25) plants along the nutrient gradient. The two pea lines are shown in different symbols, but they were not statistically different (Table S1, S2). The colours represent the nutrient levels on a log scale. The data show that most of the allocation differences represent a leaf – root trade-off along the nutrient gradient.

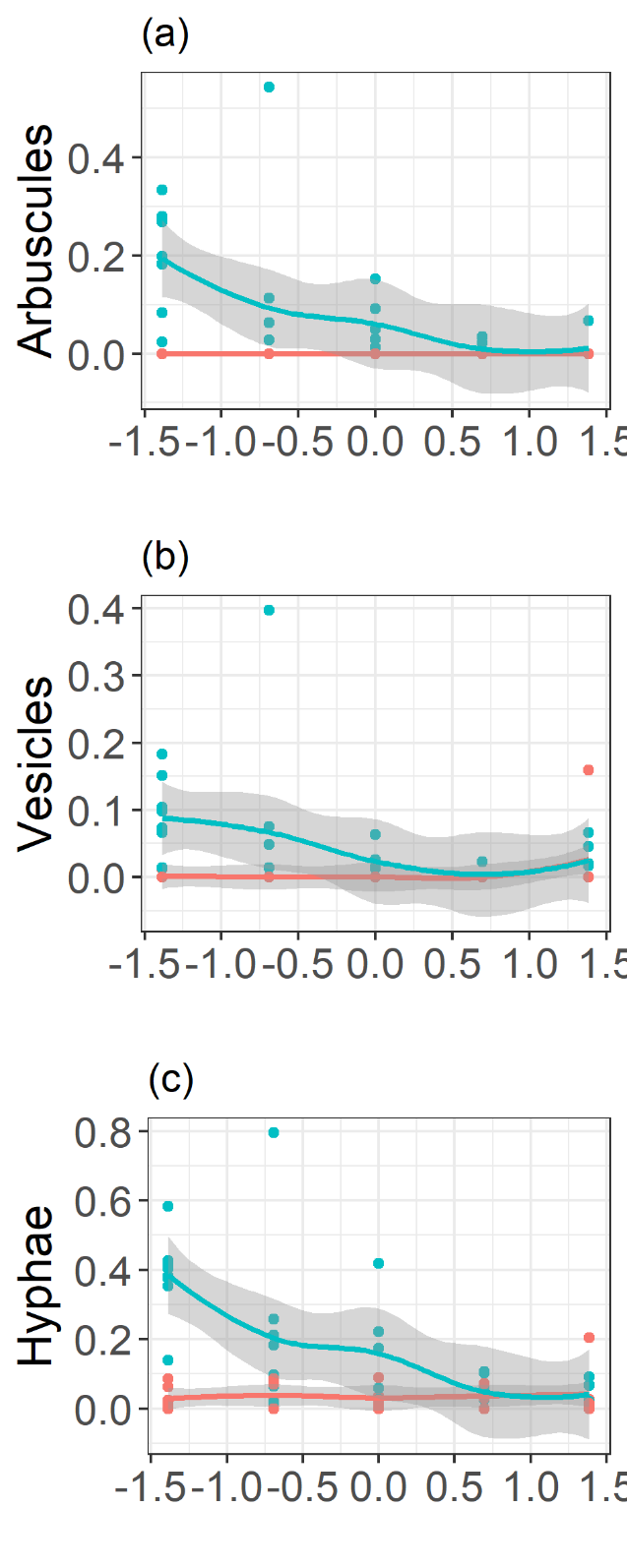

**FIG S3:** Proportion of root length colonised by mycorrhizal structures along the nutrient gradient. Wild type plants are shown in blue, mutant plants in red. Structures shown are (a) arbuscules, (b) vesicles and (c) intra-radicular hyphae. Lines represent loess regressions, and the grey bands are standard errors.

| **TABLE S1:** Results of GLMM (general linear mixed effects models) on plant growth in two pea lines (wild type and R25) using the lmer and lmerTest libraries in R. Denominator degrees of freedom (Den df) were estimated using the Satterthwaite approximation. Nutrients were treated as a continuous variable on a log scale. Nutrients were manipulated on a log scale and were analyzed on that scale. Biomass data were log transformed for normality. The model included block as a random effect. | | | | |
| --- | --- | --- | --- | --- |
| **Shoot** | | | | |
| **Factor** | **Num df** | **Den df** | **F** | **p** |
| log(Nutrients) | 1 | 65.53 | 51.13 | **<0.0001*** |
| Genotype | 1 | 66.31 | 0.06 | 0.0624 |
| Log(Nutrients) X Genotype | 1 | 65.65 | 0.51 | 0.5089 |
| **Root** | | | | |
| **Factor** | **Num df** | **Den df** | **F** | **p** |
| log(Nutrients) | 1 | 69.00 | 35.09 | **<0.0001*** |
| Genotype | 1 | 69.00 | 0.58 | 0.4503 |
| Log(Nutrients) X Genotype | 1 | 69.00 | 0.02 | 0.9013 |
| **Fruit** | | | | |
| **Factor** | **Num df** | **Den df** | **F** | **p** |
| log(Nutrients) | 1 | 66.68 | 5.21 | **0.0257*** |
| Genotype | 1 | 67.22 | 5.50 | **0.0219*** |
| Log(Nutrients) X Genotype | 1 | 66.68 | 0.61 | 0.4366 |

| **TABLE S2:** Results of GLMM on plant relative allocation to vegetative tissues in two pea lines (wildtype and R25) using the lmer and lmerTest libraries in R. Denominator degrees of freedom were estimated using the Satterthwaite approximation. Nutrients concentrations were manipulated on a log scale and were analysed on that scale. Relative allocation was arcsine square root transformed for continuity. The model included block as a random effect. | | | | |
| --- | --- | --- | --- | --- |
| **Proportion Leaf** | | | | |
| **Factor** | **Num df** | **Den df** | **F** | **p** |
| log(Nutrients) | 1 | 65.67 | 12.81 | 0.0006 |
| Genotype | 1 | 66.2 | 0.75 | 0.3889 |
| Log(Nutrients) X Genotype | 1 | 65.76 | 0.02 | 0.8940 |
| **Proportion Stem** | | | | |
| **Factor** | **Num df** | **Den df** | **F** | **P** |
| log(Nutrients) | 1 | 65.20 | 368.82 | <0.0001 |
| Genotype | 1 | 65.50 | 3.022 | 0.0868 |
| Log(Nutrients) X Genotype | 1 | 65.24 | 0.36 | 0.54967 |
| **Proportion Root** | | | | |
| **Factor** | **Num df** | **Den df** | **F** | **P** |
| log(Nutrients) | 1 | 65.27 | 311.73 | <0.0001 |
| Genotype | 1 | 65.58 | 1.38 | 0.2447 |
| Log(Nutrients) X Genotype | 1 | 65.32 | 0.25 | 0.6170 |

| **TABLE S3:** Results of GLMM on AMF colonisation using the lmer and lmerTest libraries in R. Denominator degrees of freedom were estimated using the Satterthwaite approximation. Proportion data were arcsine transformed for continuity and normality. The model included block as a random effect. | | | | |
| --- | --- | --- | --- | --- |
| **Arbuscules** | | | | |
| **Factor** | **Num df** | **Den df** | **F** | **p** |
| log(Nutrients) | 1 | 59.14 | 24.09 | <0.0001 |
| Genotype | 1 | 60.51 | 44.35 | <0.0001 |
| Log(Nutrients) X Genotype | 1 | 59.81 | 26.70 | <0.0001 |
| **Vesicles** | | | | |
| **Factor** | **Num df** | **Den df** | **F** | **p** |
| log(Nutrients) | 1 | 59.87 | 3.79 | 0.0564 |
| Genotype | 1 | 62.32 | 22.73 | <0.0001 |
| Log(Nutrients) X Genotype | 1 | 61.07 | 8.86 | 0.0042 |
| **Intraradicular Hyphae** | | | | |
| **Factor** | **Num df** | **Den df** | **F** | **P** |
| log(Nutrients) | 1 | 59.28 | 20.16 | <0.0001 |
| Genotype | 1 | 61.33 | 26.06 | <0.0001 |
| Log(Nutrients) X Genotype | 1 | 60.28 | 21.07 | <0.0001 |

**REFERENCES**.

Chapin, F.S., Vitousek, P.M. & Vancleve, K. (1986) The nature of nutrient limitation in plant-communities. *American Naturalist,* **127,** 48-58.

Johnson, N.C., Graham, J.H. & Smith, F.A. (1997) Functioning of mycorrhizal associations along the mutualism-parasitism continuum. *New Phytologist,* **135,** 575-586.

Mcgonigle, T.P., Miller, M.H., Evans, D.G., Fairchild, G.L. & Swan, J.A. (1990) A new method which gives an objective-measure of colonization of roots by vesicular arbuscular mycorrhizal fungi. *New Phytologist,* **115,** 495-501.

Mcnickle, G.G., Gonzalez-Meler, M.A., Lynch, D.J., Baltzer, J.L. & Brown, J.S. (2016) The world's biomes and primary production as a triple tragedy of the commons foraging game played among plants. *Proceedings of the Royal Society B: Biological Sciences,* **283,** 20161993.
